# De novo design of allosterically controlled enzymes

**DOI:** 10.64898/2026.09.08.750245

**Authors:** Alexander Shida, Kelly Wang, Hojae Choi, Samuel Pellock, Adam Broerman, Saman Salike, William Grubbe, Emily Joyce, Alex Kang, Asim K. Bera, Xinting Li, Arvind Pillai, David Baker

## Abstract

The ability of enzymes to sense signals and respond by changing their structure and activity underlies cellular processes from signaling to metabolic control. While there have been recent advances in the de novo design of enzymes and conformationally switching proteins, combining these to achieve allosteric regulation of enzymatic activity in a fully designed system remains an outstanding challenge. Here, we show that denoising diffusion models enable the design of compact de novo enzymes whose activity can be allosterically activated or repressed by designed protein effectors. Our results establish a general route to allosteric control over a wide range of catalytic and other protein functions.

## Intro

Virtually every cellular process relies on the ability of enzymes to respond to signals by altering their activity. Cell signaling cascades ^1^, metabolic pathways^2,3^, and motor proteins ^4,5^ often rely on coupling between enzymatic activity and protein-ligand binding at sites distal to the active site. The exact mechanisms by which native allosteric enzyme systems function are still somewhat unclear; some models suggest that sidechain communication networks relay changes from an allosteric binding site to the enzyme active site^6,7^, or that allosteric effectors act more broadly to shift the equilibria between well-defined distinct states^8–10^. Efforts to design allosteric systems in synthetic biology have generally relied on natural enzymes engineered with topological rearrangements^11,12^, insertions/fusions^13^, and split chains ^14,15^ to couple a protein-ligand binding event to enzymatic activity. Although switchable enzymes can be generated by this approach, the coupled functions are limited to natural enzyme activities, and such domain rearrangements do not structurally recapitulate the allostery in many naturally evolved enzyme folds, which are often compact single-domain proteins. Recently, there has been progress in the de novo design of enzymes ^16–18^ and the allosteric control of protein interactions ^19–21^, but the de novo design of allosteric enzymes remains an outstanding challenge.

We set out to use deep learning design methods to de novo design allosteric enzymes whose activity is modulated by a designed protein effector. In contrast to previous approaches ^19–21^ to the design of synthetic allosteric switches, which relied almost exclusively on the modular fusion of domains and large interdomain movements, we aimed to more closely emulate natural allosteric control, where a compact structure switches between subtly different conformations that differ in catalytic activity. We aimed to design allosteric enzymes that populate two distinct conformational states toggled by binding a designed effector: an active state, where the catalytic residues are optimally positioned to mediate the reaction, and an inactive state, where these residues deviate from this catalytically optimal geometry. In addition to the closer parallel with natural allosteric enzymes, we reasoned this design strategy should enable faster switching between states, as the energy barriers to conversion between very closely related states, where only a few residue-residue contacts must be broken or made, may in general be smaller than between very different states.

## Results

### Design approach

We developed a general design strategy to craft allosteric protein energy landscapes (Figure 1A). Starting from an active designed enzyme (Fig 1A, “Active”), we generate an alternative inactive state by partial RFdiffusion (Fig 1B). In partial RFdiffusion, a starting structure is subjected to 1-3Å of random Gaussian noise, and then denoised using RFdiffusion to generate alternative conformations; partial diffusion has had success in identifying subtly modified backbones with increased binding affinity ^22,23^, and we reasoned the conformational changes for allosteric regulation could be similarly subtle. We further design allosteric effectors that bind to either the active or inactive state and modulates the energy landscape. We design both allosteric activation systems in which the unbound enzyme has lower energy in the inactive state (Fig 1A, top left), and effector binding favors the active state, and allosteric inactivation systems (Fig 1A, top right) where effector binding conversely favors the inactive state. For the effector-dependent activator design, we used RFDiffusion to generate binders to the active enzyme state (Figure 1B, top trajectory). For the effector-dependent inactivator designs, we used RFDiffusion to generate effectors to inactive enzyme states (Fig 1B, bottom trajectory). We then used tied ProteinMPNN to generate sequences compatible with the backbones of the enzyme–effector complex as well as the isolated enzyme and effector states ^19^.

**Figure 1.**
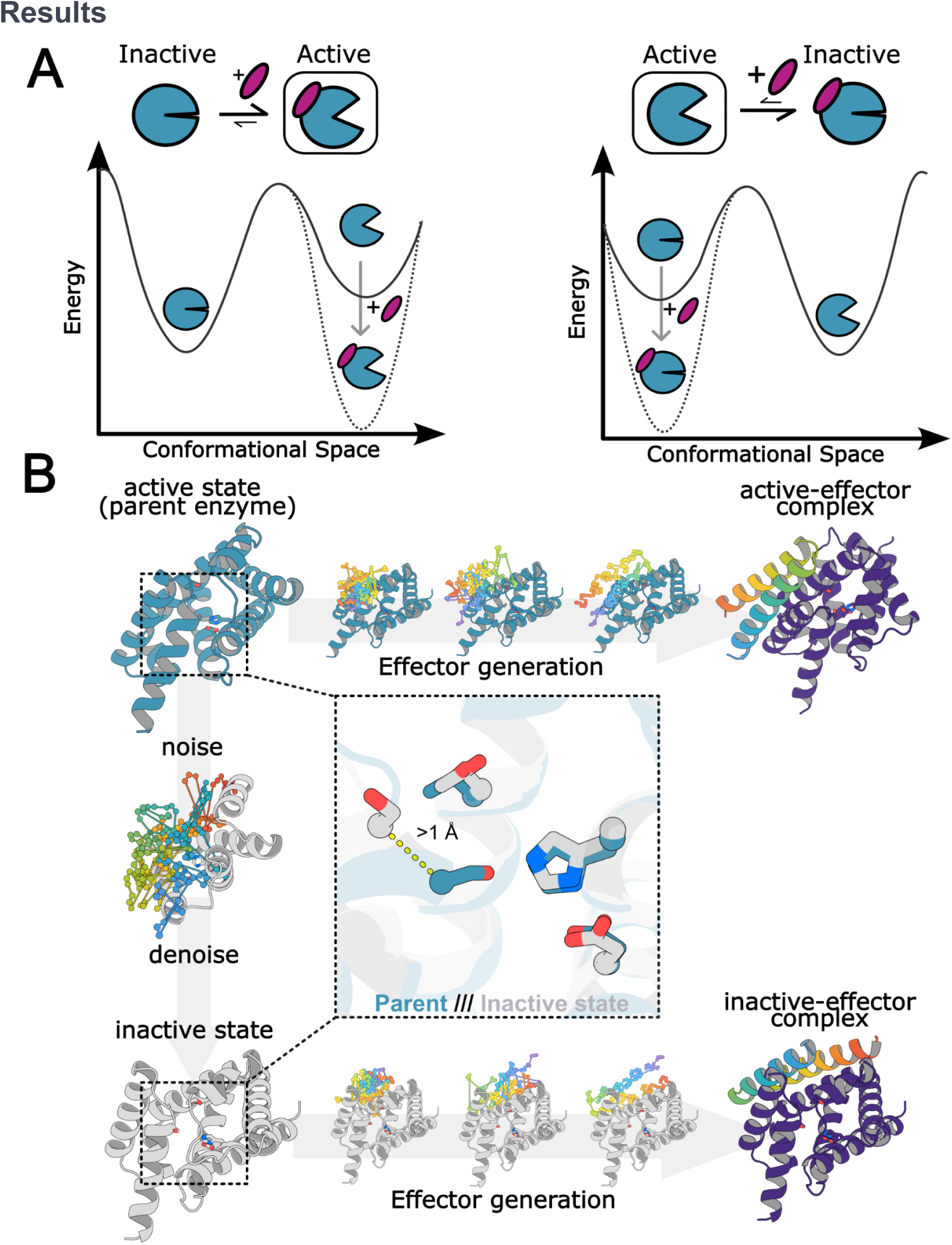
Design approach. (a) Design concept. The allosteric designed enzyme (blue circle) has two states, “Active” and “Inactive”. Binding of a designed effector (magenta oval) shifts the relative energies of the two states. In the allosteric activator case (left), the unbound enzyme has lower energy in the inactive state (solid line in left energy diagram), and binding of the effector (dotted line) stabilizes the active state. In the allosteric inhibitor case (right), effector binding leads to conversion from the active state to the inactive state. (b) Design strategy. A designed enzyme (blue, top left) is partially diffused (downward arrow) by noising and denoising a portion of the protein (colored in the middle panel at the left, the gray portion is kept fixed) to generate an alternative state (gray, bottom left) with a slightly distorted active site (inset in center, the catalytic serine has shifted position by more than an Angstrom). RFdiffusion is then used to generate binders to either the active state (top) in the allosteric activator case, or the inactive state (allosteric inhibitor case) (right panels). Sequence design using tiedMPNN is used to favor the inactive state in the absence of effector in the activator case, and the active state in the absence of effector in the inhibitor case.

We chose the *de novo* serine hydrolase Win1 as a model system as the catalytic mechanism and active-site geometry are well understood ^16^ and the design is small and compact, facilitating stabilization of alternative states. Given the precise geometry required for catalysis ^16,17^, we anticipated that the vast majority of partially diffused conformations are likely to have compromised active sites with reduced or ablated catalytic activity. To keep the energy of the perturbed state close to that of the active state to enable efficient toggling by effector, we divided the structure into two separate domains: one containing the His–Asp (catalytic dyad) and the other the catalytic Ser and oxyanion Thr (Figure 1B). We then partial diffused one of the domains and filtered for Cα displacements greater than 1 Å from the starting active-state structure (Figure 1B, left panel). Designs predicted by AlphaFold2 to adopt the target active and inactive conformations in the presence and absence of effector, respectively, were selected for experimental characterization.

### Characterization of allosteric activators

We first sought to design enzymes whose activity is allosterically upregulated upon addition of effector. As described above and outlined in Fig 1, we generated inactive win1 conformations by partial diffusion in the absence of substrate (Figure 1B) and designed small proteins that bind to and stabilize the active state of win1. Sixty-one such designed enzyme-effector pairs that passed AF2 filters were selected for experimental characterization.

The designed enzymes and effectors were expressed in *E. coli*, and activity was measured in cell lysates using the fluorogenic substrate 4-methylumbelliferone acetate (4Mu-Ac) (Figure 2A). We identified two effector-enzyme pairs, Janus1 and Janus2, for which addition of effector cell lysate substantially increased formation of the fluorescent product, 4Mu, and confirmed the enhancement of activity upon effector binding with purified proteins (Fig 2B and Fig 2D, respectively). For both pairs, there is substantial predicted conformational remodeling of the designed enzyme in the inactive and effector-bound complexes with Cα RMSDs of 8.3 Å and 3.80 Å for Janus1 and Janus2 respectively (Figure 2B and 2E, top panels). In the inactive state, the catalytic residues Ser142, His17, Asp37, and Thr99 have a disrupted geometry incompatible with efficient catalysis. Upon effector binding, both designs transition to an active-like catalytic arrangement, suggesting that effector-induced stabilization of the catalytic geometry underlies the observed increase in enzymatic activity (Figure 2B and 2E, bottom panels).

**Figure 2.**
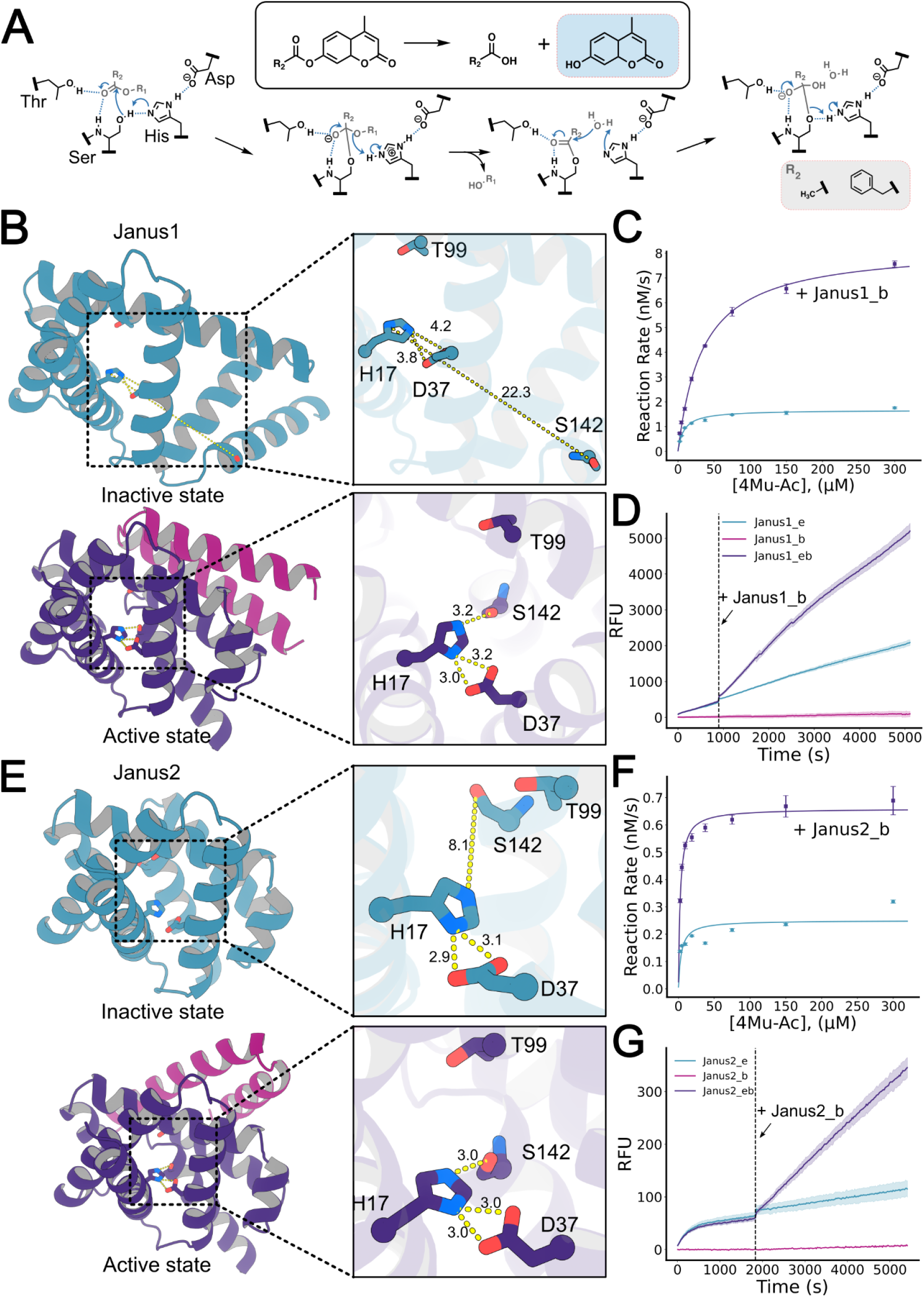
Design of allosterically activated serine hydrolases. (A) Serine hydrolase mechanism and chemical reaction. The hydrolase separates the fluorescent group, R1 in blue, from the leaving group R2 (acetate or phenylacetate). (B) Inactive and active states with catalytic active site geometry differences. (C) Michaelis-Menten kinetics of Janus1 comparing unbound enzyme and enzyme with saturating effector (D) Addition of saturating Janus1_b effector rapidly accelerates catalysis. (E-G) Same as (B-D) but with the Janus 2 system.

We characterized the mechanism of allosteric activation using mutational and kinetic analyses of the purified effectors and enzymes. Active site residue knockouts had activity reduced to background levels or below the detection limit (Supplementary Figure 1), supporting catalysis through the expected serine/cysteine hydrolase machinery, as for the parent enzyme (^16^). To characterize the kinetics in more detail, steady-state kinetic measurements were performed in the absence and presence of effector at a 1:2 enzyme:effector ratio. For Janus1, effector binding increased the turnover rate from *k*_cat_ = 0.0033 ± 0.00010 s⁻¹ to 0.017 ± 0.0003 s⁻¹, an approximately 5-fold increase (Figure 2C, Table 1). For Janus2, *k*_cat_ increased from 0.00050 ± 0.00005 s⁻¹ in the absence of effector to 0.0013 ± 0.00003 s⁻¹ in the presence of effector, corresponding to a 2.6-fold enhancement (Figure 2F, Table 1). Effector binding also altered substrate affinity in Janus1: K_m_ increased 4.3-fold, from 8.1 ± 1.0 μM in the apo state to 35 ± 2 μM in complex with Janus1_b (Figure 2C, Table 1). In contrast, the K_m_ of Janus2 was unchanged within error (4 ± 2 μM apo vs. 2.5 ± 0.3 μM bound; Figure 2F, Table 1).

**Table 1.**
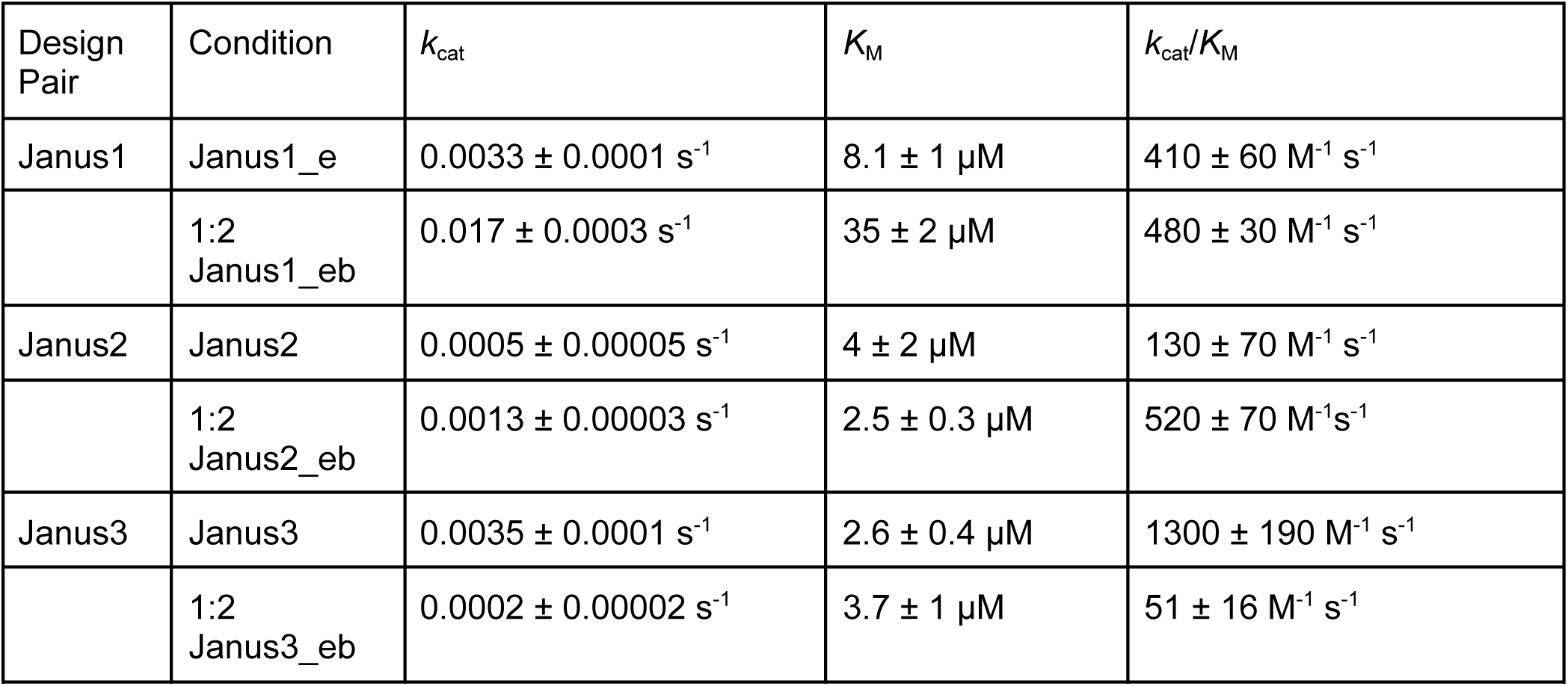

| Design Pair | Condition | $k_{\text{cat}}$ | $K_{\text{M}}$ | $k_{\text{cat}}/K_{\text{M}}$ |
| --- | --- | --- | --- | --- |
| Janus1 | Janus1_e | $0.0033 \pm 0.0001 \text{ s}^{-1}$ | $8.1 \pm 1 \text{ }\mu\text{M}$ | $410 \pm 60 \text{ M}^{-1} \text{ s}^{-1}$ |
| | 1:2<br>Janus1_eb | $0.017 \pm 0.0003 \text{ s}^{-1}$ | $35 \pm 2 \text{ }\mu\text{M}$ | $480 \pm 30 \text{ M}^{-1} \text{ s}^{-1}$ |
| Janus2 | Janus2 | $0.0005 \pm 0.00005 \text{ s}^{-1}$ | $4 \pm 2 \text{ }\mu\text{M}$ | $130 \pm 70 \text{ M}^{-1} \text{ s}^{-1}$ |
| | 1:2<br>Janus2_eb | $0.0013 \pm 0.00003 \text{ s}^{-1}$ | $2.5 \pm 0.3 \text{ }\mu\text{M}$ | $520 \pm 70 \text{ M}^{-1} \text{ s}^{-1}$ |
| Janus3 | Janus3 | $0.0035 \pm 0.0001 \text{ s}^{-1}$ | $2.6 \pm 0.4 \text{ }\mu\text{M}$ | $1300 \pm 190 \text{ M}^{-1} \text{ s}^{-1}$ |
| | 1:2<br>Janus3_eb | $0.0002 \pm 0.00002 \text{ s}^{-1}$ | $3.7 \pm 1 \text{ }\mu\text{M}$ | $51 \pm 16 \text{ M}^{-1} \text{ s}^{-1}$ |

To estimate the rate of onset of allosteric activation with enzyme engaged in active catalysis, we performed spike-in experiments in which a 2x molar ratio of effector was added to an ongoing enzymatic reaction. For both Janus1 and Janus2, addition of effector increased catalytic activity to the levels observed in the pre-incubation experiment. The onset of activation was too rapid to resolve a lag phase under our assay conditions, indicating that the effector on-rate is fast relative to the mixing and measurement dead time (∼21 seconds; Figure 2D and 2G, bottom right panels). The cognate effectors for Janus1 and Janus2 rapidly bind to and shift their target enzymes into a more catalytically competent active state. To test the specificity of each effector for its cognate enzyme, we incubated Janus1 and Janus2 with Janus2_b and Janus1_b, respectively. Both enzymes demonstrate specificity to their respective effector, with no rate enhancement when incubated with the non-cognate effector (Supplementary Figure 2).

### Characterization of allosteric inhibitors

We used a similar approach to design allosteric inactivators. We based these designs off of designed serine hydrolases, win1_b1 ^16^ and win1_b4 (Supplementary Figure 1), that hydrolyze the larger substrate 4-methylumbelliferone phenylacetate (4Mu-PhAc, Figure 2A). Inspired by the coupling in molecular machines like Hsp70 ^24^, in which ATP binding switches Hsp70 into a protein-binding competent state, we reasoned that the bulkier, more hydrophobic leaving group of 4Mu-PhAc would provide more binding energy to drive conformational change. During initial experimental characterization we observed reduced catalytic activity relative to the parent enzyme. To expand the dynamic range for measurement and more readily enable observation of allosteric suppression of activity, we sought to improve the baseline catalytic activity of the designed enzymes in the unbound form. We found that a serine-to-cysteine substitution led to dramatic increases in catalytic efficiency across a variety of designed serine hydrolases (Supplementary Figure 1,4). This cysteine mutation was included in all subsequent designs.

We experimentally characterized 24 designed allosteric inhibitor-enzyme pairs using lysate screening with fluorescent 4Mu-PhAc in the absence and presence of the effector (Supplementary Figure 3). Of these, eight designs retained catalytic activity, and one, Janus3, had a clear reduction of enzymatic activity for both cysteine and serine variants upon addition of effector. Structural comparison of the predicted active and effector-bound inactive-state complexes shows more subtle conformational remodeling in Janus3 compared to Janus1 and Janus2, with Cα RMSDs between states of 1.84 Å (Figure 3A). In the active state, the catalytic residues Cys142, His17, Asp37, and Thr99 adopt a geometry consistent with the catalytically active parent enzyme. Upon effector binding, the nucleophilic Cys142 is disrupted and turned outward and away from the catalytic pocket.

**Figure 3.**
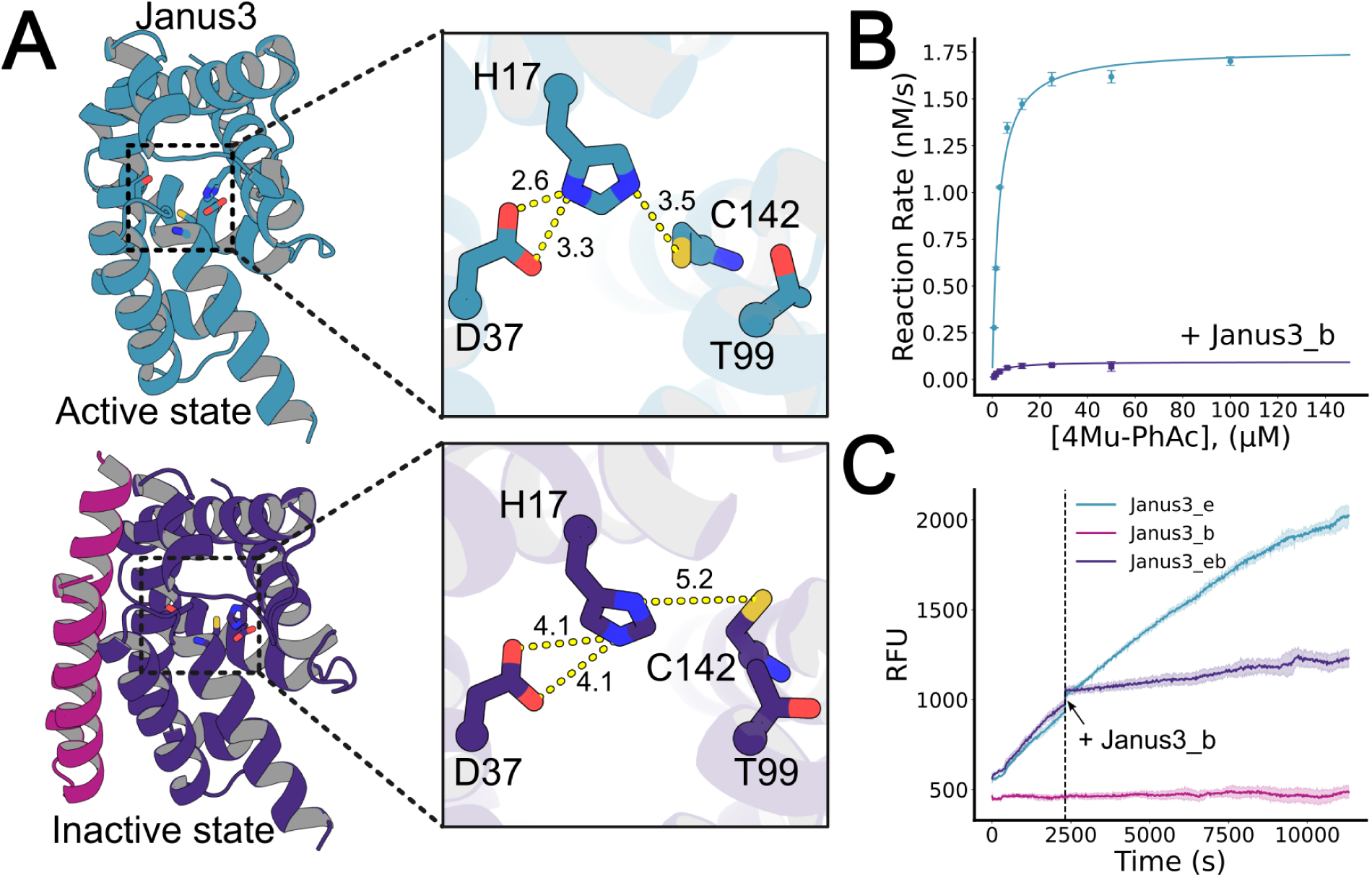
Characterization of the Janus3 allosteric inhibition system. (A) Design models of the active and inactive states of Janus3 with close-ups of the active site changes. The primary change is in the catalytic nucleophile residue (B) Michaelis-Menten kinetics of Janus3 comparing unbound enzyme and enzyme with saturating effector. (C) Addition of saturating effector while reaction is in progress rapidly reduces catalytic rate.

We characterized the allosteric inhibition mechanism of the Janus3 enzyme and effector (Janus3_b) with proteins purified from *E.coli*. Measurement of rate as a function of substrate concentration revealed a substantial decrease in *k*_cat_ upon Janus3_b binding (Figure 3B) from .0024 s⁻¹ (unbound) to .0003 s⁻¹ (bound), an 8-fold reduction (Table 1). There is no significant difference in *K*_M_ between the bound and unbound states of Janus3, suggesting that the binding of Janus3_b alters catalytic residue geometry without affecting the substrate binding pocket of Janus3, consistent with the design model. Mutation of active site residues Cys142, His17, Asp37, and Thr99 to alanine eliminated or reduced activity, confirming that the catalytic mechanism is unchanged (Supplementary Figure 1). In spike-in experiments, upon addition of Janus3_b into the Janus3_e and 4Mu-PhAc reaction, the rate decreased to a lower steady state within the dead time of our mixing procedure (Figure 3b, bottom right), again indicating that our designed allosteric regulation mechanism is fast on this time scale.

We succeeded in obtaining a crystal structure of the Janus3 enzyme-effector complex at 1.7 Å resolution (Figure 4A). The interface between the enzyme and effector is recapitulated precisely as designed, with a Cα RMSD of [0.77] Å to the design model over the interface residues(Figure 4A, bottom). The effector binds far from the active site, confirming that the alteration in activity occurs purely due to conformational coupling (rather than steric interference with substrate binding). The catalytic cysteine was oxidized to sulfinic acid, which likely occurred during crystallization or data collection. Beyond this modification, the geometry of the catalytic site is similar to that of the bound inactive state design model (crystal-inactive state design model Cα RMSD= 1.66 Å, crystal-active state design model Cα RMSD= 1.87 Å), and to the closely related active state design model (Figure 4B). The primary differences between the designed active vs inactive states lie in a shift in the placement of the cysteine residue and a rotation and translation of the Thr99 oxyanion stabilizing residue (Figure 4B). Both the crystal and the inactive state design model show a 1.2 Å shift of the cysteine backbone Cα relative to the active state model in addition to the rotation of the oxyanion stabilizing oxygen away from the catalytic pocket. These subtle changes show that small shifts in the active site can translate into large changes in activity, and confirm the ability of our design strategy to yield atomically accurate allosteric modulation of *de novo* enzymes.

**Figure 4.**
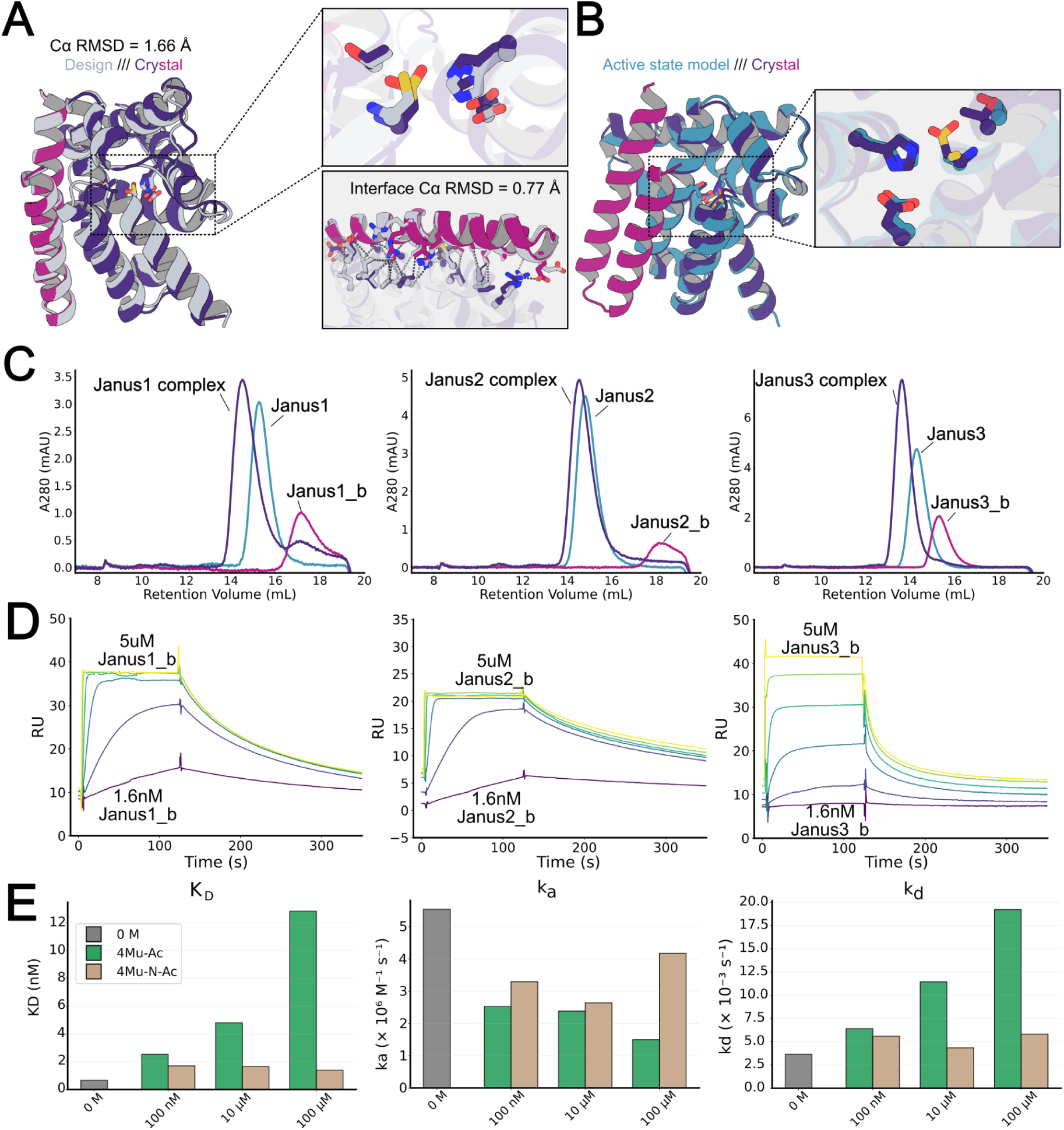
Structure and substrate modulation of effector-enzyme interface. (A) Crystal structure of Janus3 in complex with cognate effector Janus3_b. The crystal structure is overlaid on the Janus3 enzyme-effector complex at left; insets show more detailed comparisons of the catalytic residues (top inset) and interface residues (bottom inset) (B) Comparison of the crystal structure of Janus3—Janus3b complex with the active state design model of unbound Janus3 (C) SEC traces for the Janus1, Janus2, and Janus3 systems with enzyme alone, effector alone, and enzyme and effector in a 1:1 stoichiometric ratio. (D) Kinetics of association of designed effector-enzyme systems using SPR. (E) Influence of substrate and inhibitor on Janus1 effector binding. Grey zero M bar is with buffer and effector only, green is with the presence of substrate 4Mu-Ac and brown is in the presence of inhibitor 4Mu-N-Ac.

Having examined the influence of effector binding on catalysis, we next investigated the effect of substrate on effector binding. We first characterized binding of effector and enzyme in the absence of substrate for the three Janus systems. Size exclusion chromatography (SEC) of effector alone, enzyme alone, and enzyme and effector together revealed monodisperse 1:1 binding (Figure 4B); Janus2 enzyme alone appears to be a dimer, but forms a monodisperse complex with Janus2_b with the expected molecular weight (Figure 4B, middle). For all three systems, the nearly complete shift in population to the monodisperse complex peak upon addition of effector indicates near stoichiometric binding under these conditions.

We measured the binding affinity of the designed effectors for their cognate enzymes by surface plasmon resonance (SPR), immobilizing the enzymes on the sensor surface via streptavidin-biotin conjugation and flowing each cognate effector at concentrations from 5 µM to 1.6 nM in 5-fold serial dilutions (Figure 4B). All of the allosteric effectors bind their cognate enzymes with nanomolar to sub-nanomolar affinity: equilibrium dissociation constants (K_D_) of 2.97 nM for Janus1 (k_on_ = 2.52 × 10⁶ M⁻¹s⁻¹, k_off_ = 7.46 × 10⁻³ s⁻¹), 0.726 nM for Janus2 (k_on_ = 3.49 × 10⁶ M⁻¹s⁻¹, k_off_ = 2.53 × 10⁻³ s⁻¹), and 106 nM for Janus3 (k_on_ = 5.93 × 10⁵ M⁻¹s⁻¹, k_off_ = 6.31 × 10⁻² s⁻¹) (Table 2). These on-rates exceed those of previously designed de novo conformationally changing, allosteric proteins (Praetorius et al. 2023; Pillai et al. 2024; Broerman et al. 2025). The Janus3 on-rate is roughly an order of magnitude lower than those of Janus1 and Janus2, suggesting lower occupancy of the inactive binding-competent conformation and higher occupancy of the unbound, active state.

**Table 2.**

| Design | Substrate | Concentration (μM) | $K_D$ (nM) | $k_{on}$ ($M^{-1}s^{-1}$) | $k_{off}$ ($s^{-1}$) |
| --- | --- | --- | --- | --- | --- |
| Janus1 (A1) | | 0 | 2.97 | $2.52 \times 10^6$ | $7.46 \times 10^{-3}$ |
| | 4Mu-Ac | 100 | 13.3 | $1.18 \times 10^6$ | $1.58 \times 10^{-2}$ |
| | 4Mu-Ac | 10 | 4.23 | $2.09 \times 10^6$ | $8.83 \times 10^{-3}$ |
| | 4Mu-Ac | 0.1 | 4.03 | $1.85 \times 10^6$ | $7.45 \times 10^{-3}$ |
| | 4Mu-N-Ac | 100 | 2.73 | $2.86 \times 10^6$ | $7.79 \times 10^{-3}$ |
| | 4Mu-N-Ac | 10 | 1.98 | $2.12 \times 10^6$ | $4.20 \times 10^{-3}$ |
| | 4Mu-N-Ac | 0.1 | 3.14 | $2.23 \times 10^6$ | $7.00 \times 10^{-3}$ |
| Janus2 (A12) | | 0 | 0.726 | $3.49 \times 10^6$ | $2.53 \times 10^{-3}$ |
| | 4Mu-Ac | 100 | 2.34 | $3.45 \times 10^6$ | $8.08 \times 10^{-3}$ |
| | 4Mu-Ac | 10 | 0.845 | $3.29 \times 10^6$ | $2.78 \times 10^{-3}$ |
| Janus3 (A9) | | 0 | 106 | $5.93 \times 10^5$ | $6.31 \times 10^{-2}$ |
| | 4Mu-PhAc | 100 | 105 | $3.48 \times 10^5$ | $3.64 \times 10^{-2}$ |

We next used SPR to measure the binding of the effector to the enzyme at different substrate concentrations (Table 2). While there are many allosteric enzymes for which catalysis does not impact allosteric effector binding under physiologically relevant conditions (^25,26^), such coupling is a hallmark of natural molecular machines (^27–29^. For both the Janus2 and Janus3 systems, we observed no significant changes in binding constants between presence or absence of substrate concentrations (Table 2). At high substrate concentrations, we observed a modest but reproducible (Figure 4C) change in the binding affinity of the Janus1 system, with K_D_ shifting from 2.97 nM (no substrate) to 13.3 nM (with 100 µM substrate, Figure 4C, Table 2). Michaelis-Menten kinetic analyses indicated a five-fold increase in the *K*_M_ of Janus1 in the bound state, while in Janus2 and Janus3 systems the *K*_M_ values remain approximately the same (Table 1).

To determine whether the coupling between catalysis and binding for Janus 1 reflects substrate binding or catalytic turnover, we repeated the experiment using a non-hydrolyzable substrate analog containing an amide bond in place of the scissile ester linkage (Supplementary Figure 5). This analog did not change effector binding affinity relative to the substrate-free condition, with all affinities remaining near 2–3 nM (Figure 4C, Table 2). Because the nearly isosteric amide analog has no effect on effector binding across concentrations, the affinity changes with the cleavable substrate likely arise from catalytic turnover rather than substrate occupancy of the active site or off-target binding at the effector interface. Indeed, we found that the covalent acyl enzyme intermediate accumulates during active catalysis (Supplementary Figure 8); this covalent species is likely more able to outcompete binding of the much larger protein effector than the non-covalently interacting inhibitor.

## Discussion/Conclusion

We demonstrate that allosteric enzymes can be designed by using generative AI methods to perturbed an enzyme and designing effectors that specifically stabilize either this state (allosteric inhibitors) or the original active state (allosteric activators). We find that allosteric modulation occurs rapidly (within the dead time of our mixing experiments, and with high on-rates), as in natural systems, but distinct from a previously designed allostery design system in which large scale conformational changes were required (^20^). These results suggest that for design of rapidly switching allosteric systems, smaller magnitude conformational changes can have lower energy barriers to reconfiguration. Our partial diffusion approach broadly explores the local conformational space of a given protein and can achieve subtle forms of allosteric modulation that resemble natural systems and were previously difficult to attain. The close agreement of the crystal structure of the Janus3 designed effector-enzxyme complex with the design model demonstrates the atomic accuracy of our allostery design approach. There is still room for improvement in the structure prediction methods used to predict state in the presence and absence of effector, in the calculation of the free energy difference between the two states, and in the method of generation of the alternative state; in the latter case multistate design methods should in principle be able to directly optimize for an inactive state in the absence and the active state in the presence of a co-designed effector.

Because our designs resemble native allosteric systems in both the magnitude of the conformational change and the rapidity of response, the designed mechanism may inform our understanding of natural allostery. It has been suggested that sidechain communication pathways connect allosteric sites with active sites, enabling transmission of information between them ^6,7^. However, we do not explicitly design any such communication networks; instead we focus entirely on design of two distinct states whose energy differences are toggled by effector–the presumption is that the two are in rapid equilibrium and the effector functions by modulating the equilibrium between the two of them. Our approach for designing allosterically controlled enzymes should be adaptable to a wide variety of exciting applications; for example in metabolic engineering enzymes that carry out key steps in pathways for generating new high value compounds, where the extent of activity can be, for example, made dependent on the abundance of substrate (in the activator case) or product (in the inhibitor case). Allosteric de novo enzymes could also provide an effective route to new classes of sensors, where a protein analyte is sensed through its allosteric effect on the activity of an easy to monitor enzyme, complementing current approaches relying on domain insertion ^11^.

A particularly interesting application of allostery is to the design of machines and motors. If effector binding was modulated through the catalytic cycle, then hydrolysis of a fuel molecule could be coupled to the binding and unbinding of the foot of a walker from a track. We anticipated at the outset of our design efforts that the effects of effector on catalysis and of catalysis on effector binding may be reciprocal as they are thermodynamically coupled. Overall, we observed stronger effects of effector binding on catalysis than and weaker effects of substrate binding on effector binding; catalysis is likely more sensitive to small sub-Angstrom changes in conformation, and hence the effects on k_cat_ are much larger than the perturbation of binding of substrate on the effector, or the perturbation of binding of effector on substrate. We do observe a change in effector binding affinity for Janus1 over the full catalytic cycle which likely reflects formation of the acyl enzyme intermediate; in natural systems such as the Ras family of small GTPases, protein binding affinities are similarly modulated throughout the catalytic cycle ^30^. More generally, the design of efficient machines and motors coupling a fuel cycle with protein binding and unbinding will require design of stronger coupling between substrate binding, catalysis and effector binding, an exciting next challenge for protein design.

## Methods

### Computational design of allosteric enzymes

#### Partial RFDiffusion for inactive state generation

We used existing *de novo* serine hydrolase enzymes from Lauko et al. as our starting active enzyme backbone ^16^. The ligand substrate was removed from the design model and partial diffusion (T = 20-30) was run in separate batches on residues 88-158, 1-87, and 1-158 to generate alternative backbone conformers (Figure 1).

#### RFDiffusion for effector generation

Small protein effectors (30-80 residues) for either the original, active state or the newly generated, inactive states of the enzyme were generated using RFDiffusion. The `ppi.hotspot_res` parameter localized the interface of the effector generation to residues distal to the catalytic active site of enzyme (‘A100’, ‘A103’, ‘A104’, ‘A107’, ‘A110’, ‘A125’, ‘A128’, ‘A132’, ‘A135’, ‘A136’). The `inpaint_seq` flag was used for the entire protein (A1-158) to allow for more proximal effector placement. The resultant effectors were designed to be either allosteric activators or inhibitors depending on whether they were generated in the context of the active enzyme state or inactive enzyme state, respectively.

#### Backbone filtering

The generated inactive enzyme states from the partial diffusion step were first filtered for topological consistency by computing DSSP assignments for both the original, active state and each of the partially diffused states. Second, structural similarity was quantified by performing a Ca backbone alignment of the structures based on the specific residue ranges that were fixed during the partial diffusion step. For the structures generated without fixing any residues, a global Ca alignment was performed. Distance based measurements along the key catalytic residues of the enzymes were used to select only generated states that had significant (>1 A) movement in one of the catalytic Ca atoms.

The effector backbone designs were individually scored for compactness and contacts with the enzyme. Designs with contact scores greater than 0.2 and compactness scores greater than 0 were accepted.

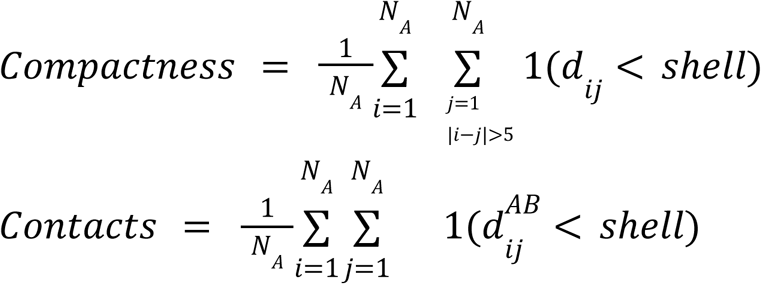

Where N_A_ = number of residues in chain A, dij = distance between CA atoms of residues i and j in chain A, and 1(⋅) = indicator function (1 if true, 0 if false). The condition |i - j| > 5 excludes local sequential neighbors and *shell* was set to 6 angstroms. 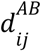 is the distance between CA atom of residue i in chain A and CA atom of residue j in chain B

#### Sequence design

The allosteric system designs were composed of the active and inactive enzyme states, and an effector in complex with one of the two states for allosteric activation or inhibition, respectively. We then used ProteinMPNN to generate sequences for each allosteric activator grouping (two sequences. one for both enzyme states and one for the effector). We leveraged the `symmetry_residues` and `symmetry_weights` flags to enforce residue correspondence despite the conformational differences between the inactive and active states.

#### Design filtering

We then used AF2 to predict the structure for each sequence with and without the presence of its cognate effector. Designs were selected based on their correspondence to the initial generated backbone designs as well as quality of predicted fold (RMSD < 2; mean_pae < 5; pLDDT > 85). For the allosterically inhibited enzyme systems, we additionally used AlphaFold3 to model the AF2-passing designs in two states: an effector-bound state (enzyme, effector, and substrate) and an unbound state (enzyme and substrate without effector). We created a structural filter designed to assess whether the predicted ground-state complex was geometrically competent for the first chemical step, nucleophilic attack of the Ser142 Oγ on the scissile ester carbonyl carbon with an intact His–Asp charge-relay. For each model, active-site geometry was quantified from the predicted coordinates of the catalytic triad and the bound substrate: the His17 ND1–Asp37 carboxylate distances (the His–Asp leg of the relay), the His17 NE2–Ser142 Oγ distance (positioning of His to activate the nucleophile), the Ser142 Oγ–carbonyl carbon distance and attack angle (Oγ···C–O, measured at the carbonyl carbon), and the relative His–Ser orientation captured by two angles and a dihedral. Model confidence was summarized by the complex pLDDT.

Each model was assigned an “active” score by counting how many criteria consistent with a catalytically competent triad it satisfied: complex pLDDT > 85; both His ND1–Asp OD1 and ND1–Asp OD2 distances < 3.3 Å; His NE2–Ser Oγ distance < 4.0 Å; Ser Oγ–carbonyl carbon distance < 3.85 Å; and the His–Ser top angle (64–95°), bottom angle (74–104°), Oγ–carbonyl attack angle (70–100°), and His–Ser dihedral (−115 to −72°) each falling within its target window. Each satisfied criterion contributed +1. A complementary “inactive” score counted criteria indicating a disrupted active site—His ND1–Asp distances > 4 Å, His NE2–Ser Oγ distance > 4.0 Å, and Ser Oγ–carbonyl carbon distances exceeding 5, 7, and 9 Å (scored cumulatively)—with each satisfied criterion contributing −1. A per-state score was defined as the sum of the active and inactive scores.

To prioritize designs predicted to function as effector-induced inactivators, designs were ranked by the difference between the unbound and bound per-state scores (score_delta = unbound − bound) in descending order. Designs at the top of this ranking were those whose active-site geometry was predicted to be competent for nucleophilic attack in the unbound state and disrupted upon effector binding. This ranking was used to order candidates for subsequent expert review; final designs were selected by expert visual inspection of active-site geometry.

### Screening of modulated enzymatic activity

#### DNA cloning, expression and crude purification of designs

DNA sequences for the designed proteins were synthesized as eBlocks from IDT and inserted into the pCOOL1 vector (addgene) using GoldenGate cloning^16^. This vector includes an mScarlet-i3 tag fused to the C-terminus, allowing activity measurements in lysate to be normalized by enzyme concentration. The resulting plasmids were introduced into NEB BL21(DE3) cells, which were then grown in 1 ml overnight cultures within 96-well deep-well round-bottom plates (2 ml capacity) at 37°C and 1300 rpm on a Heidolph shaker. For protein production, 50 μl aliquots from overnight cultures were transferred to 1 ml autoinduction media in 96-well deep-well round-bottom plates (2 ml capacity) and incubated for roughly 24 hours at 37°C and 1300 rpm. Cells were pelleted by centrifugation (4000g, 10 min), the supernatant was removed, and pellets were washed with buffer (20 mM HEPES, 50 mM NaCl, pH 7.4) by resuspending on the Heidolph shaker for 5 min at room temperature and 1300 rpm. After another centrifugation step (4000g, 10 min) and supernatant removal, pellets were lysed by resuspension in 500 μl lysis buffer (20 mM HEPES, 50 mM NaCl, 0.01 mg/ml DNAse I, 0.01 mg/ml lysozyme, 1 mM EDTA, 0.1% Triton X-100) and incubated for 2 hours at 37°C and 1300 rpm. Lysates were clarified by centrifugation (4300g, 30 min) and the soluble fraction was collected for assays. Activity assays were performed by dispensing 3 μl of the enzyme lysate with 3uL of either buffer (20 mM HEPES, 50 mM NaCl, pH 7.4) or 3 ul of its cognate effector lysate into microtiter plates and starting reactions with 54 μl of substrate buffer (111.1 μM 4MU-Ac or 4MU-PhAc in 20 mM HEPES, 50 mM NaCl, pH 7.4, 5% DMSO). After substrate addition, plates were first read for mScarlet-i3 fluorescence, then continuously monitored for 4MU product formation (ex: 365 nm, em: 445 nm) using a Neo2 plate reader.

### Characterization of identified hits

#### DNA cloning, expression, and purification of allosteric enzymes

Successful design pairs were selected for further analysis. Designs were purified with either a C-terminal 6xHis tag, a StrepTactin tag (www.iba-lifesciences.com), or a C-terminal AviTag followed by a 6xHis tag for subsequent site specific biotinylation. All proteins were expressed in NEB BL21(DE3) *E. coli* using TBII (MpBio) autoinduction media with 0.5% (w/v) glycerol, 0.05% (w/v) glucose, 0.2% (w/v) lactose, 20 mM MgSO4, trace metal mix and 50 μg ml−1 kanamycin. Expression cultures (50mL) were grown at 37°C overnight, collected by centrifugation, resuspended and lysed in 15 mL of lysis buffer (20 mM HEPES, 50mM NaCl, 0.5 mg/mL polymyxin B, 0.1 mg/mL lysozyme, 0.01 mg/mL DNAse), and then clarified with centrifugation at 14,000g for 45 minutes. Clarified lysate containing soluble protein was then bound to 1ml Ni-NTA resin (QIAGEN) or 1mL Strep-Tactin XT 4Flow resin (IBA Lifesciences), washed with either high salt wash buffer (500 mM NaCl, 40 mM imidazole, 100 mM KPO4, pH 7.4) or Strep-Tactin wash buffer (100 mM Tris, 150 mM NaCl, 1mM EDTA, pH 8.0), and eluted with Ni-NTA elution buffer (500 mM imidazole, 40 mM Tris, 300 mM NaCl) or StrepTactin Elution buffer BXT (IBA Lifesciences, 100 mM Tris, 150 mM NaCl, 1 mM EDTA, 50 mM biotin), respectively. The elution containing our protein was then further purified using SEC on a fast protein liquid chromatography (FPLC) system (AKTA Pure) with a Superdex 75 increase 10/300 GL column in a HEPES-buffered solution (20mM HEPES, 50mM NaCl, pH 7.4) into 1-mL fractions. Fractions corresponding to monomers and expected elution fractions were collected and concentration was measured using the estimated molar extinction coefficients predicted from protein sequences and integrating the absorbance at 280 nm over the collected fractions. Collected proteins were then analyzed on liquid chromatography-mass spectrometry to confirm correct molecular weights corresponding to the designed constructs.

#### Kinetics assay

To characterize hits identified from lysate screens for catalytic turnover, we incubated purified protein samples with fluorogenic substrates 4Mu-Ac and 4Mu-PhAc. Kinetic screens were performed in 40 μl reaction volumes in 96-well half-area plates, 60 μl reaction volumes in 96-well full-area plates, or 20 μl reaction volumes in 384-well plates. Protein and substrate were prepared fresh in 20 mM HEPES, 50 mM NaCl, pH 7.4, 5% DMSO. Enzyme was added to microtiter plates at a final concentration of 500 nM, and for effector-bound conditions effector was included at a 2:1 molar ratio relative to enzyme. Reactions were initiated by addition of substrate, applied as a two-fold serial dilution series starting from 150 μM or 100 μM. Generation of the fluorogenic product 4Mu was monitored continuously (excitation 365 nm, emission 445 nm) on a Neo2 plate reader with incubation at 30°C. Analysis of the resulting data was carried out using custom scripts (see computational methods). Raw fluorescence values were converted to 4Mu product concentrations using a linear standard curve, and signal wells were blank-corrected against substrate-only wells. Initial velocities were determined as the slope of a linear fit to the blank-corrected product progress curve over an early time window exhibiting a steady-state rate. Initial velocities from replicate wells at each substrate concentration were averaged (mean ± SEM), and the resulting velocity-versus-[S] data were fit to the Michaelis-Menten equation by non-linear least squares to obtain V_max_ and K_m_. k_cat_ was calculated as V_max_/[E] using the enzyme concentration in the assay, and k_cat_/K_m_ was derived accordingly; parameter uncertainties were taken from the covariance matrix of the fit and propagated to k_cat_ and k_cat_/K_m_. To determine the uncatalyzed reaction rate in assay buffer (20 mM HEPES, 50 mM NaCl, pH 7.4, 5% DMSO), substrate was diluted in buffer alone and rates determined at multiple substrate concentrations, after which the rate was determined from fitting [S] versus rate with an equation of the form rate = k_buffer_[S].

#### SEC binding assay

To validate that allosteric enzyme-effector pairs were forming complexes we measured shifts in the SEC profiles of each design pair with and without their designed partner. For each design pair three conditions were prepared: enzyme alone, effector alone, or enzyme and effector together. The purified proteins were diluted to a final concentration of 5uM in 0.5 or 1 mL volumes and injected serially using an automated FPLC system (AKTA Pure) with a flow rate of 0.5 mL min^-1^ on a Superdex Increase 75 10/300 GL column. Absorbance signals at 230 and 280 nm were measured to monitor the elution profile of protein across the run.

#### SPR assay

Allosteric enzymes to be captured on the SPR chip were first purified with a C-terminal AviTag and subsequently biotinylated with 5 ug ml-1 BirA (Avidity), 10 mM ATP, 10 mM Mg(OAc)2 and 100 uM D-biotin and incubated at room temperature for at least 4 hours before further purification by SEC. Successful purification was confirmed using LC-MS. SPR measurements were performed at 25°C in HBS-EP+ buffer (Cytiva) with 5% DMSO on a Biacore 8K instrument. Biotinylated proteins were immobilized on the chip using the Biotin CAPture system (Cytiva).

For measurements of rate constants between the allosteric effector and enzyme, first the chip was equilibrated with modified HBS-EP+ buffer with 5% DMSO and varying concentrations of small molecule substrate (4Mu-PhAc or 4Mu-Ph-N-Ac; 0M, 100uM, 10uM, 100 nM) was flowed over the chip for 150s, following this, allosteric effector was flowed over the chip in an association step of 120 s, finally the same modified HBS-EP+ buffer containing substrate used during equilibration is flowed over chip to attain dissociation rates. Each channel of the SPR went through multiple cycles of these three steps for varying concentrations of effector (5uM, 1uM, 200nM, 40nM, and 8nM) while maintaining the same concentration of small molecule.

#### Crystallography

Crystallization experiments were conducted using the sitting drop vapor diffusion method. Initial crystallization trials were performed via sitting-drop vapor diffusion in 96-well plates at 20 °C, using a mosquito LCP liquid handler (SPT Labtech) to dispense 200 nl drops. Plates were imaged using UVEX microscopes and UVEX PS-256 from JAN Scientific. For Janus3-Janus3_b complex structure, diffraction quality crystals formed in 0.1 M HEPES pH 7.5 and 1.4 M Sodium citrate tribasic dihydrate. 2M L-Proline used as cryo protectant during flush freezing in liquid N_2_ before sending to the synchrotron center for diffraction experiment.

Diffraction data were collected at either the National Synchrotron Light Source II (NSLS-II) beamline 17-ID-2 (FMX). X-ray intensities and data reduction were evaluated and integrated using XDS ^31^ and merged/scaled using Pointless/Aimless in the CCP4 program suite ^32^. Structure determination and refinement starting phases were obtained by molecular replacement using Phaser ^33^using the designed model for the structures. Following molecular replacement, the models were improved using phenix.autobuild ^34^ and using simulated annealing. The structure was refined in Phenix ^34^. Model building was performed using COOT ^35^. The final model was evaluated using MolProbity ^36^. Data collection and refinement statistics are recorded in the statistics table below. Atomic coordinates and structure factors reported in this paper have been deposited in the Protein Data Bank (PDB), with accession code pdb_000037UG.

