## Supplemental Information for "De novo design of allosterically controlled enzymes"

Supplementary Figure 1

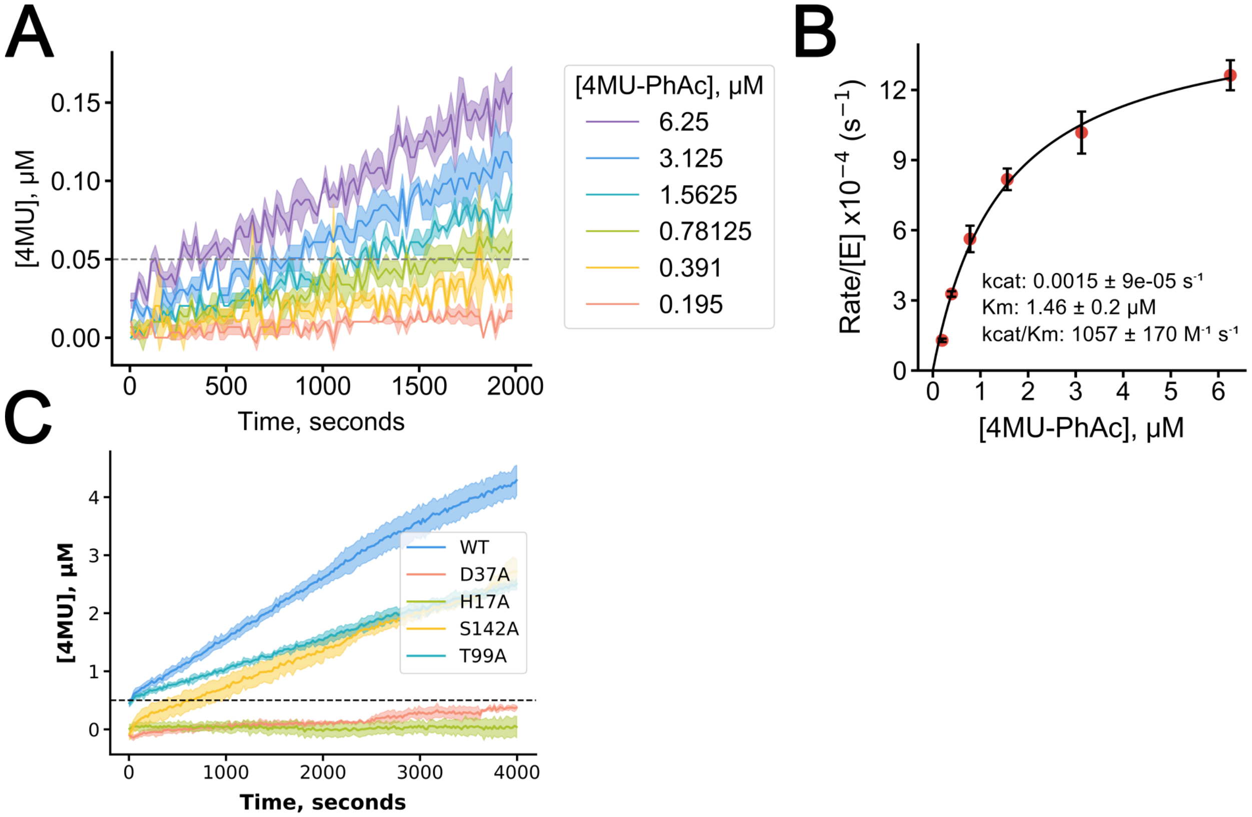


Designed serine hydrolase, win1_b4, that is catalytically active on the 4-methylumbelliferone phenylacetate (4Mu-PhAc) substrate. Designed via partial diffusion of the original win1 enzyme with 4Mu-PhAc in the active site, following the design strategy described by Lauko et al. [16] (A) Catalytic activity of 50nM of win1_b4 incubated with 4Mu-PhAc at various concentrations. (B) Michaelis-Menten kinetics of win1_b4 with a $k_{cat}$=0.0015 $s^{-1}$ and $K_M$=1.46 μM (C) Catalytic residue knockouts demonstrate reduced or ablated activity.

Supplementary Figure 2

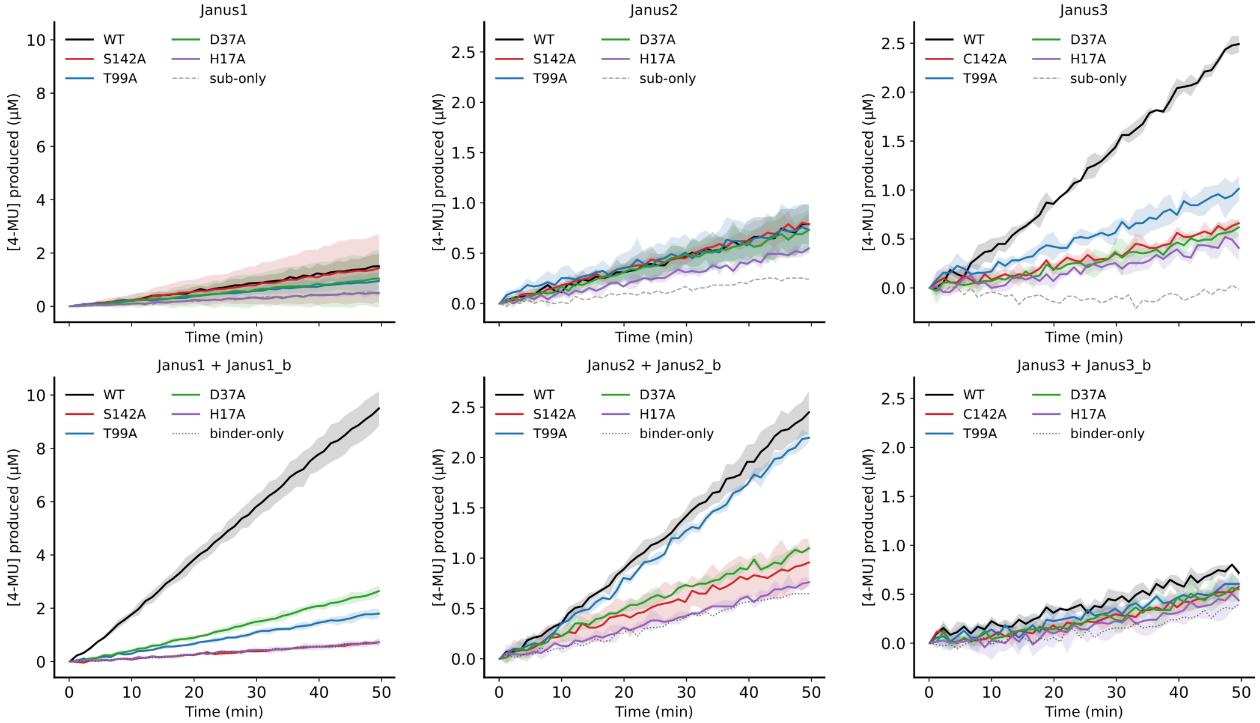


Catalytic residue knockouts for each of the allosteric enzyme systems demonstrate reduced/ablated catalytic activity. All three enzymes share the same four catalytic residues: Asp17, His37, oxyanion stabilizing Thr99, and nucleophilic Ser or Cys 142 and are mutated to Alanine.

Supplementary Figure 3

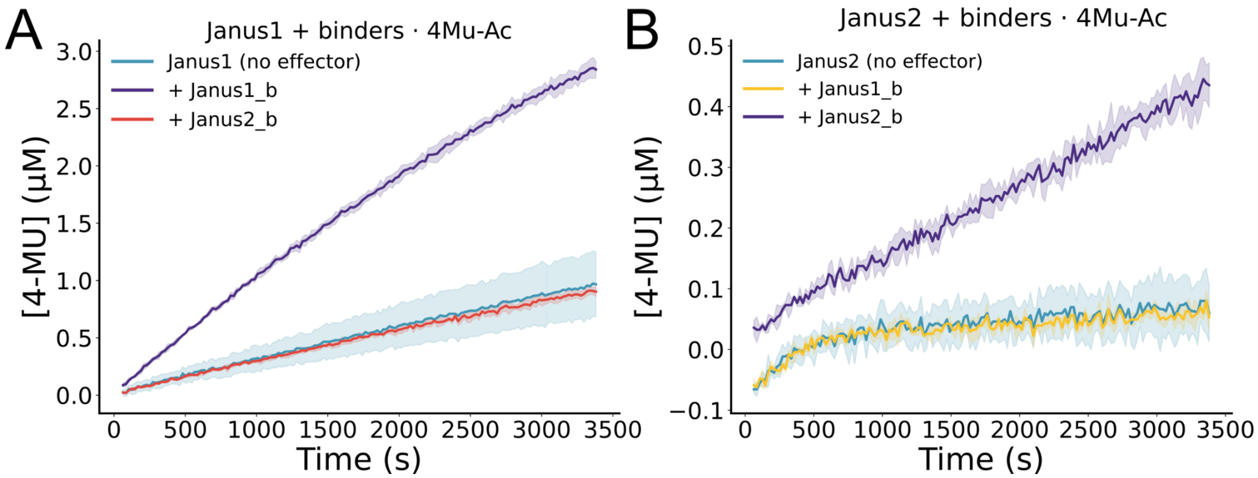


Specificity of Janus effectors. Janus1 and Janus2 hydrolysis activity on 4Mu-Ac in the presence of no effector, cognate effector, or noncognate effector. Noncognate effector conditions demonstrate no rate enhancement compared to the cognate effector.

Supplementary Figure 4

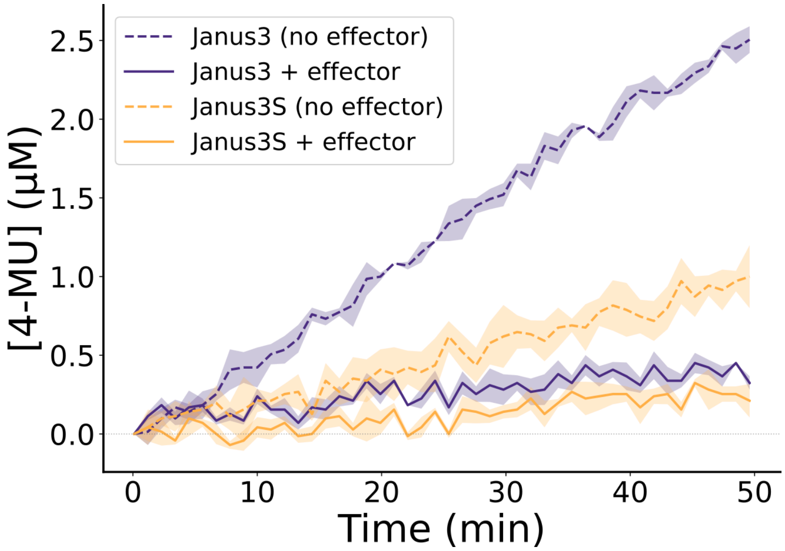


Comparison between the cysteine hydrolase Janus3 and the serine mutant (C142S) Janus3S demonstrates enhanced catalytic activity of the cysteine hydrolase. Both are shown with and without their cognate effector acting as an inhibitor. Reactions

Supplementary Figure 5

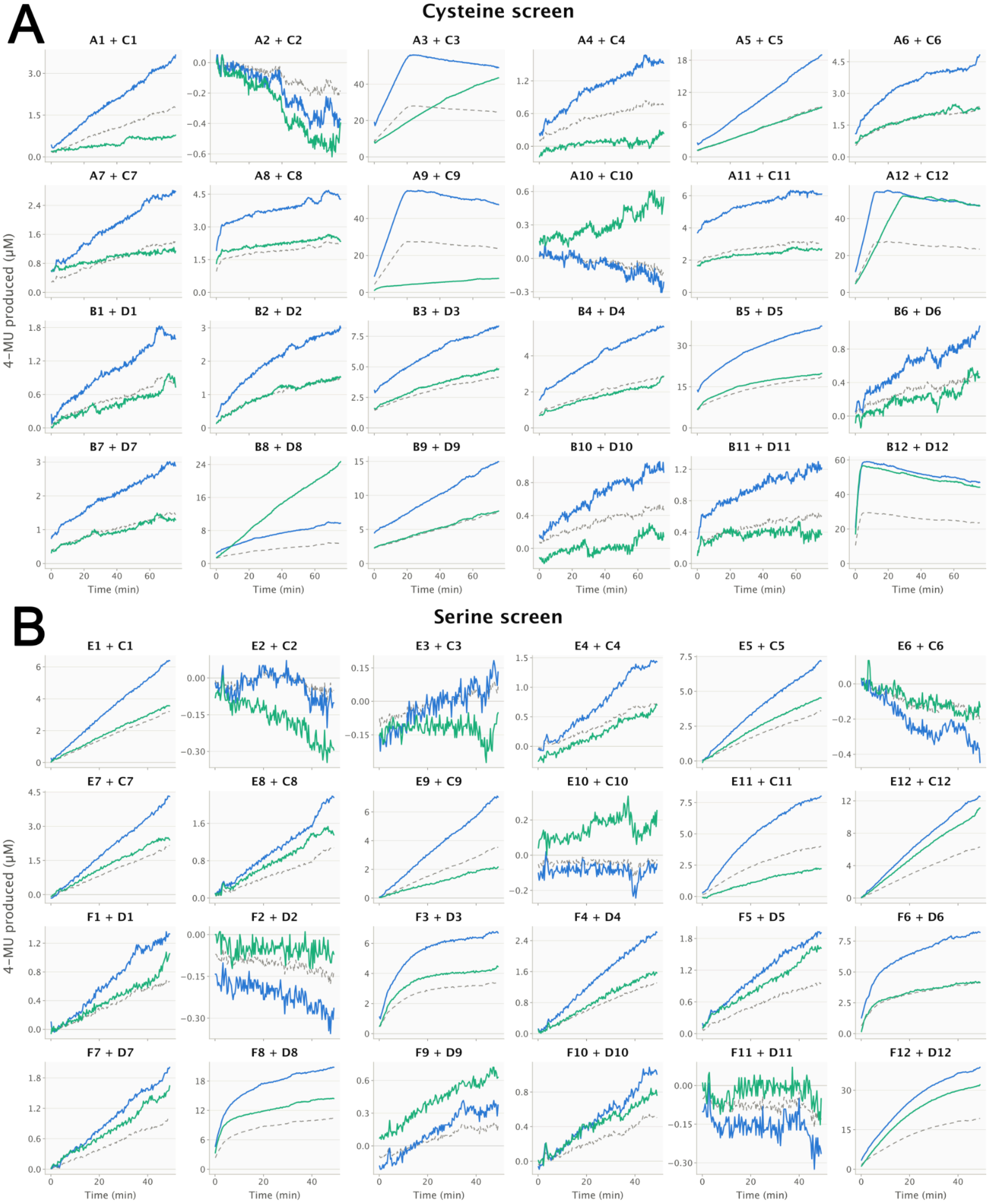


Effector dependence of 4-MU release across the cysteine (A) and serine (B) design sets. Each subpanel is one enzyme–effector pair, labelled enzyme + effector (24 pairs per set). Traces are background- and lysis-corrected as described in Methods: the buffer-plus-substrate blank and then the matched effector-only well, which carries no designed catalytic activity, were subtracted from every condition. Blue, enzyme alone; green, the 1:2 enzyme–effector mix; dashed grey, 0.5 × the enzyme-alone trace (the trajectory expected for the mix if the effector does not alter activity). A green trace falling below the dashed line therefore indicates effector-dependent suppression beyond simple dilution. n = 1 well per condition; x axes are shared within each panel (0–76 min in A, 0–49 min in B) and y axes are scaled independently per subpanel. The cysteine (A) and serine (B) designs correspond one-to-one between panels with just a single C142S mutation.

Supplementary Figure 6

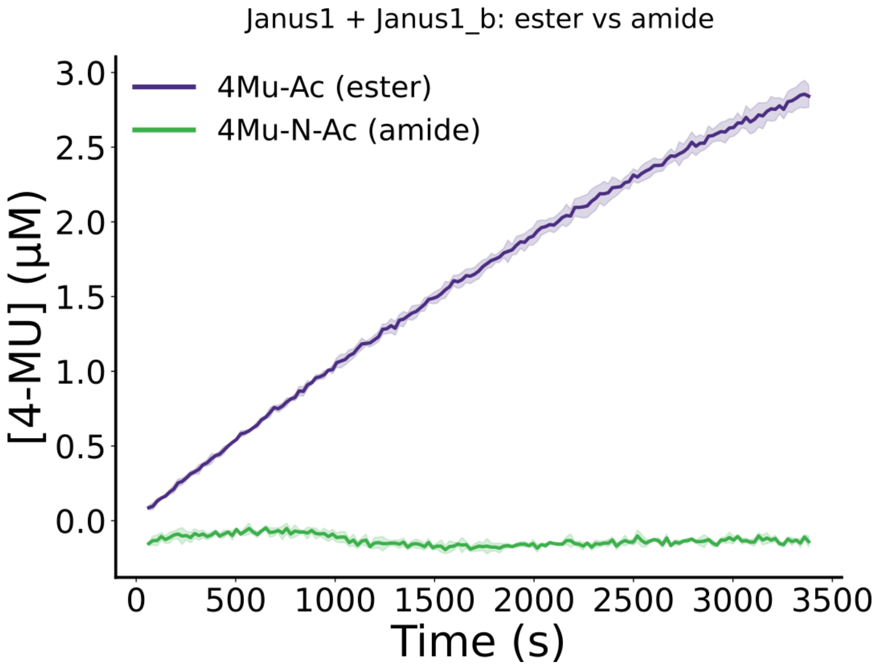


Catalytic activity of the active Janus1-Janus1_b complex showing activity on the 4Mu-Ac substrate but no activity against the amide analog substrate, 4Mu-N-Ac. 50uM of each substrate were incubated with a mixture of 1uM of Janus1 and 2uM of Janus1_b.

Supplementary Figure 7

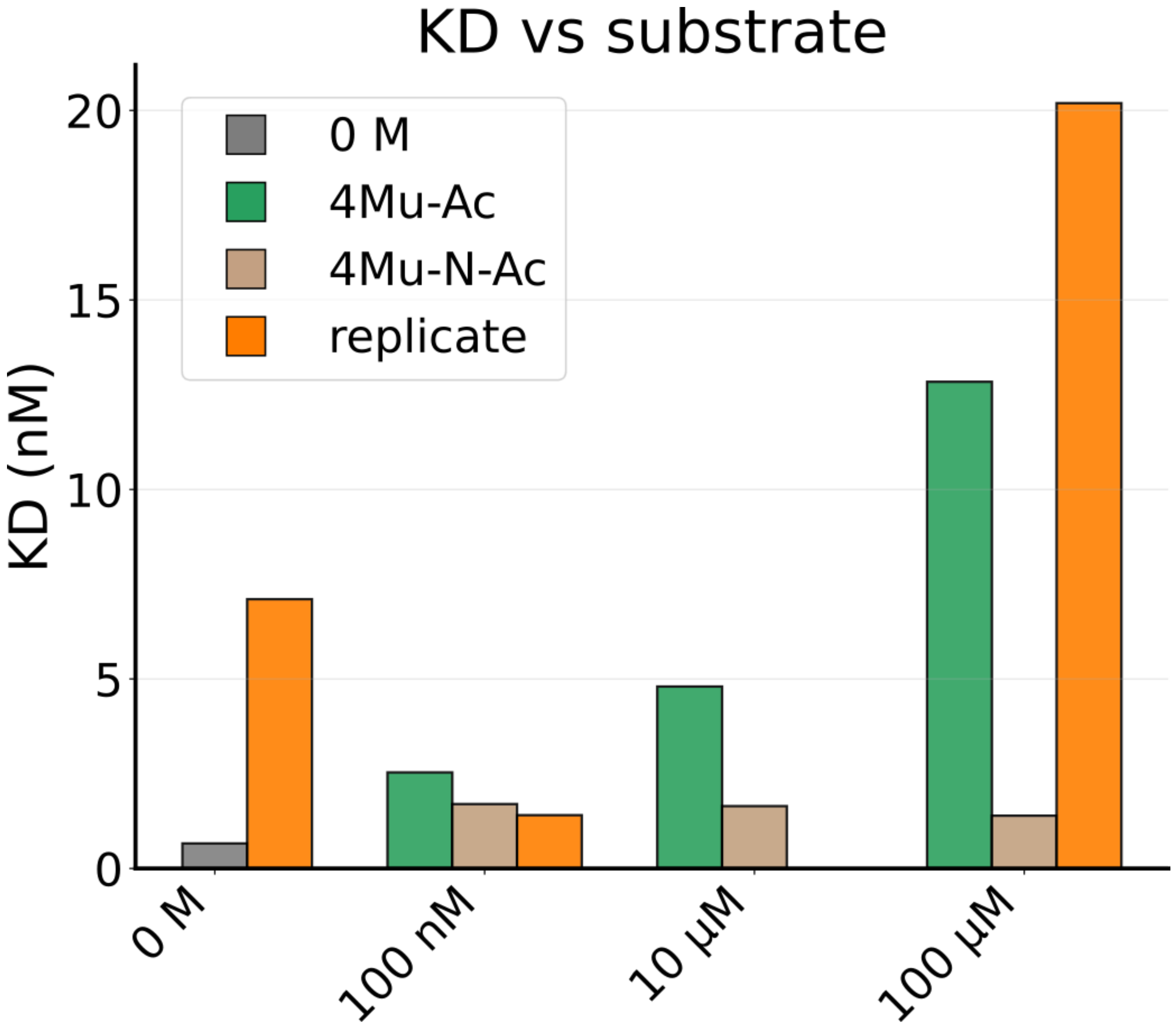


Experimental replicates of SPR experiment flowing Janus1_b over Janus1 affixed to the surface of the chip. Replicates demonstrate a consistent trend of decreased binding affinity as concentration of the cleavable substrate 4Mu-Ac is increased. Relative to the consistent $K_D$ values observed with the uncleavable 4Mu-N-Ac substrate across the same concentration regimes.

Supplementary Figure 8

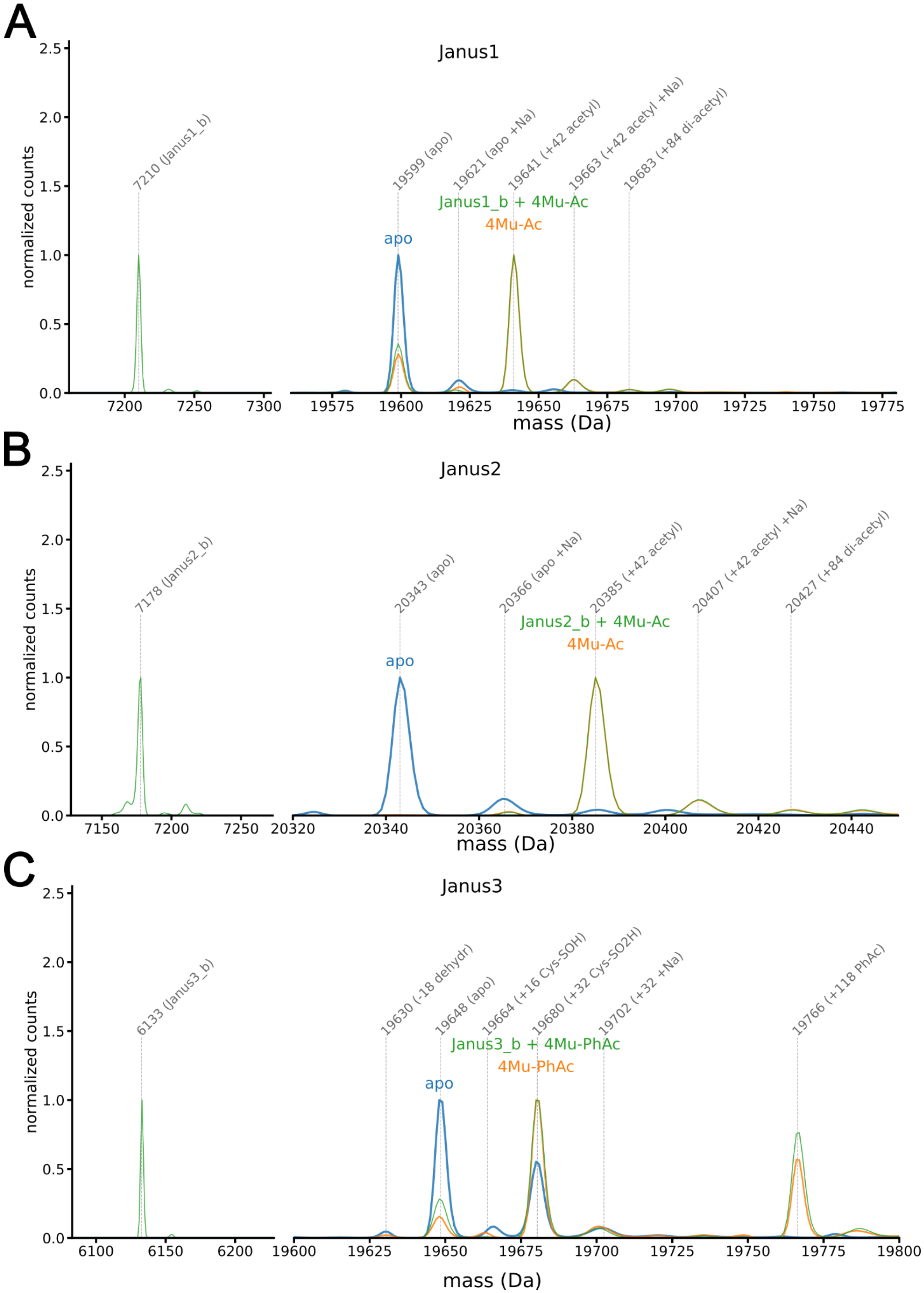


Mass spectrometry of Janus1 (A), Janus2 (B), and Janus3 (C) from 2% formic acid quenched reactions under three different conditions: apo, with their respective substrate, and with their respective substrate and cognate effector. (A) Janus1 results show expected peaks at the predicted mass of both Janus1 and Janus1b and acyl intermediates in both the bound and unbound conditions when present with 4Mu-Ac. (B) Janus2, similar to Janus1 with less variance between relative populations of each peak. (C) Janus3 shows expected peaks at predicted masses in all conditions, with additional cysteine oxidation species and a peak with the phenylacetyl enzyme intermediate.

Supplementary Table 1

| | | | | |
|---|---|---|---|---|
| Janus4 | Janus4 | 0.0011 ± 0.0002 $s^{-1}$ | 1.33 ± 1.25 μM | 829 ± 794 $M^{-1}$ $s^{-1}$ |
| | 1:2 Janus4_eb | 0.0006 ± 0.0001 $s^{-1}$ | 0.56 ± 0.51 μM | 1080 ± 993 $M^{-1}$ $s^{-1}$ |

**Crystallographic Table : Data collection and refinement statistics**

| | Janus3-Janus3_b complex **(PDB ID: pdb_00037UG)** |
|---|---|
| **Data collection** | |
| Space group | $P\ 2_1\ 2_1\ 2_1$ |
| Cell dimensions | |
| *a*, *b*, *c* (Å) | 40.49, 68.05, 71.62 |
| α, β, γ (°) | 90, 90, 90 |
| Resolution (Å) | 31.69 - 1.70 (1.74 - 1.70) |
| $R_{merge}$ | 0.15 (1.45) |
| $I / \sigma I$ | 8.0 (1.78) |
| Completeness (%) | 99.88 (100) |
| Redundancy | 13.3 (13.6) |
| $CC_{1/2}$ | 0.998 (0.728) |
| **Refinement** | |
| Resolution (Å) | 31.69 - 1.70 (1.74 - 1.70) |
| No. reflections | 22410 (1580) |
| $R_{work}$ / $R_{free}$ | 0.2293 (0.4212) /0.2791 (0.4392) |
| No. atoms | |
| Protein | 1605 |
| Ligand/ion | 1 |

| | |
|---|---|
| Water | 71 |
| *B*-factors | |
| Protein | 33 |
| Ligand/ion | 37 |
| Water | 38 |
| R.m.s. deviations | |
| Bond lengths (Å) | 0.005 |
| Bond angles (°) | 0.680 |

*Single Crystal used for each data collection. *Values in parentheses are for highest-resolution shell.


### Acknowledgments

Crystallographic diffraction data were collected at the National Synchrotron Light Source II (NSLS-II). This work used beamline 17-ID-2 (FMX), supported by the Center for BioMolecular Structure (CBMS). CBMS is primarily supported by the NIH/NIGMS through a Center Core P30 Grant (P30GM133893) and by the DOE Office of Biological and Environmental Research (KP1605010). NSLS-II is a U.S. DOE Office of Science User Facility operated for the DOE Office of Science by Brookhaven National Laboratory under Contract No. DE-SC0012704.
